# Low-flux electron diffraction study on the intercellular lipid organization in the human lip stratum corneum

**DOI:** 10.1101/2023.09.01.555969

**Authors:** Junko Kamimoto-Kuroki, Hiromitsu Nakazawa, Hiroki Ohnari, Mutsumi Yamanoi, Eiji Naru, Satoru Kato

## Abstract

**OBJECTIVE:** Chapped lips, characterized by drying and cracking, remain a prevalent concern. Identifying the root causes of lip chapping is crucial for developing effective treatments. We examined the lateral packing structure of intercellular lipids (ICL) in the lip stratum corneum (SC) by low-flux electron diffraction (LFED) to obtain new insights into the causes of high transepidermal water loss (TEWL) and low water retention, which may increase the vulnerability of the lip to chapping.

**METHODS:** Twenty-one healthy subjects participated in this study. After water content and TEWL measurements, a layer of corneocytes was collected from each lip vermilion surface by the grid-stripping technique. The lateral packing structure of ICL on the collected corneocytes was analyzed by LFED.

**RESULTS:** Similar to skin SC-ICLs, we found coexistence of orthorhombic and hexagonal phases in lip SC-ICLs. We also found that electron diffraction (ED) images with no sharp peaks and a relatively small broad peak at around 2.2 nm^−1^ appeared frequently, unlike skin SC-ICLs. This suggests that a large fraction of corneocytes in the lip SC is surrounded by thin ICL layers in the fluid phase. Such structural features of lip SC-ICLs can explain its inferior barrier function. Furthermore, we calculated the frequency of appearance of ED images with no sharp peaks, *A*_*f*_, and quantitatively analyzed its correlation with water content and TEWL. The analysis showed a negative correlation between *A*_*f*_ and water content when *A*_*f*_ > 50%.

**CONCLUSION:** This is the first report on the detailed analysis of lipid organization in lip SC-ICLs. We showed that the LFED method in combination with quasi-noninvasive sample collection by the grid-stripping technique is useful for statistical study of the fine structures in lip SC. We also found that the proportion of ICLs in a fluid phase was much higher in lip SC than in skin SC, which may be related to lower water content and vulnerability of lip to chapping. Our findings provide a promising approach for obtaining clues to the structural factors regulating the water content and TEWL in lip SC, leading to more effective lip care products.

## Introduction

The lip is critical in characterizing a viewer’s impression of our face [1]. Healthy lips can make a person’s facial expression lively, and both women and men currently use various lip care products to keep their lips healthy. The lip is thus a major research target for cosmetic and pharmaceutical researchers.

The lip is located at the boundary between the facial skin and the oral mucosa, the muco-cutaneous junction [2]. The morphological characteristics of the lip are different from those of the facial skin and oral mucosa. For example, the lip has no sweat glands or hair follicles, although these are widely distributed on facial skin. The stratum corneum (SC) is present in both the lip and facial skin but not in the oral mucosa, and lip SC is thinner than facial SC [3–5]. The presence of ceramides specific to SC supports the presence of SC in the lip, although the distribution of these ceramide species and their chain lengths differ from those in other body sites [6–8]. Functionally, the lip SC exhibits a lower water content and higher transepidermal water loss (TEWL) than cheek skin SC [9, 10]. These two parameters are frequently used as indicators of two essential SC functions: moisture-retention, and the skin barrier. Morphological and functional inadequacies of lip SC are believed to make the lip more prone to severe chapping, resulting in hard-to-cure multilayer desquamation, cracking, and bleeding. Moreover, the water content is lower in chapped lips than in healthy lips, and there is a negative correlation between lip SC hydration and the severity of chapping [11-13].

It is thus important clinically as well as cosmetically to clarify the factors causing inadequate lip SC functions, especially structural factors at the molecular level, which could shed light on how to improve SC functions pharmaceutically. However, there is essentially no molecular-level structural information on lip SC to our knowledge. For instance, orthorhombic and hexagonal phases coexist in skin SC and are known to play an important role in skin SC barrier function, but it remains unknown how widely distributed these structures are in lip SC [14, 15].

In this study, we attempted to establish a non-invasive method to analyze the lateral packing structure of lip SC-ICLs. We used a low-flux electron diffraction (LFED) method primarily because it allows us to obtain clear diffraction images of a single corneocyte collected using the grid-stripping technique with minimal invasiveness [16-20]. The LFED method is particularly suitable to fine structure analysis of lip SC compared to X-ray diffraction [14], which requires the invasive collection of a large amount of sample, or Raman spectroscopy, which presents difficulties in obtaining stable data from the narrow, soft lip region [21]. We analyzed electron diffraction (ED) images of corneocytes collected from 21 volunteers and found that both orthorhombic and hexagonal phases coexist in lip SC-ICLs, similar to skin SC-ICLs. Furthermore, we observed that the lip corneocytes surrounded by thin lipid layers in the fluid phase are extensively distributed in lip SC.

## Materials and Methods

### Subjects

Ten healthy male and 11 healthy female (total 21) subjects, aged 22 to 38 years, participated in this study. All subjects gave informed consent through both written documentation and oral communication to participate. The study was performed in accordance with the Declaration of Helsinki and approved by the Shiba Palace Clinic Ethics Review Committee (approval codes 143258_rn-26764, 144024_rn-27498 and 146407_rn-29684). Lower lip vermilions were used for all measurements after the subjects had washed their lips with a facial wash. The subjects were acclimated for 20 min in a testing room (temperature: 25 ± 2°C, humidity: 45 ± 5%) prior to sample collection. The subjects were asked not to apply lip balm or similar products to their lips for at least two weeks prior to sample collection and were forbidden to lick their lips during the experiments.

## Evaluation of lip stratum corneum functions

### Water content

Water content was measured on the lower lip vermilion using an MY-808S Skin Moisture Checker (Scalar Corp., Tokyo, Japan). A 5 mm square electrode was used to confine the measurement area within the vermilion region, which had a width of approximately 10 mm [22, 23]. Measurements were performed 5 times per subject, and the average value of 3 measurements (excluding the maximum and minimum values) was used as the estimated water content of the lip SC of the subject.

### Transepidermal water loss

TEWL was measured on the lower lip vermilion using a closed-chamber Vapometer device (Delfin Corp., Kuopio, Finland) operated in standard measurement mode. We used an attachment with an inner-diameter of 4.5 mm (Model number: SWL2040-09) instead of the standard attachment with an inner-diameter of 11 mm to fit the width of the vermilion. Measurements were performed five times per subject and the average value of three measurements (excluding the maximum and minimum values) was used as the estimated TEWL for the subject. The obtained values were corrected by multiplying by 5.98, which is the ratio of the area of the standard attachment to that of the attachment used in this experiment.

## Low-flux electron diffraction

### Sample preparation by the grid-stripping technique

A layer of corneocytes was collected from each lip vermilion surface by the grid-stripping technique for electron diffraction analysis [16-18] after water content and TEWL measurements on the same surface spot. Briefly, a thin 600 mesh bar grid (Nisshin EM Corp. Ltd., Tokyo, Japan) was coated with a glue (POLYTHICK, Sanyo Chemical Industries Ltd., Kyoto, Japan) and stored for over 24 h before use to completely remove the solvent. After stripping the uppermost layer of the lower lip vermilion once with an adhesive tape, the glue-coated grid was pressed onto the vermilion surface at several different positions to collect as many corneocytes as possible.

### Electron diffraction of corneocytes

The corneocytes collected by the grid stripping method were placed in a conventional transmission electron microscope (TEM, JEM1400, JEOL Ltd., Tokyo, Japan) operated at 100 kV, and their ED images were obtained using a high-sensitivity CCD camera (ES500W Erlangshen, Gatan, PA, USA) at room temperature. We moved the specimen more than 30 μm between shots to obtain one ED image per cell. The instrumental camera length was set at 150 cm and an electron beam was irradiated perpendicular to a single flat corneocyte adhered to the grid. The electron beam was adjusted to have a very low flux of 2.8 e·nm^−2^ s^−1^ and the exposure time was set to 0.5 s to avoid electron beam damage [16, 17]. The total electron dose of 1.4 e·nm^−2^ is low enough to suppress beam damage. A set of 10 to 25 ED images was acquired for each subject. ED intensity profiles were calculated as a function of the magnitude of the scattering vector *s* = 2sin*θ*/*λ*, where 2*θ* is the scattering angle and *λ* is the wavelength of the incident electron beam, by integrating the intensity along the azimuthal direction.

## Results and Discussion

### Electron diffraction images of cells collected from lip stratum corneum

A layer of cells was stripped from the lip surface of each of the 21 subjects (10 male and 11 female) using the grid stripping technique (Figs 1 and 2) and ED images were taken (Fig 3). The ED images and the diffraction intensity profiles were classified into three types: ED images showing (I) two sharp Debye-Scherrer rings at *s* ≈ 2.4 nm^−1^ and *s* ≈ 2.7 nm^−1^ (Figs 3 and 4 (a)), (II) one sharp Debye-Scherrer ring at *s* ≈ 2.4 nm^−1^ (Figs 3 and 4 (b)), and (III) one very broad Debye-Scherrer ring at around *s* ≈ 2.2 nm^−1^ (Figs 3 and 4 (c and d)). The most conspicuous feature of the lip specimens was the abundance of the last type-III ED images (63% of the total ED images obtained), which are rarely observed in the skin [17, 24].

**Figure 1.**
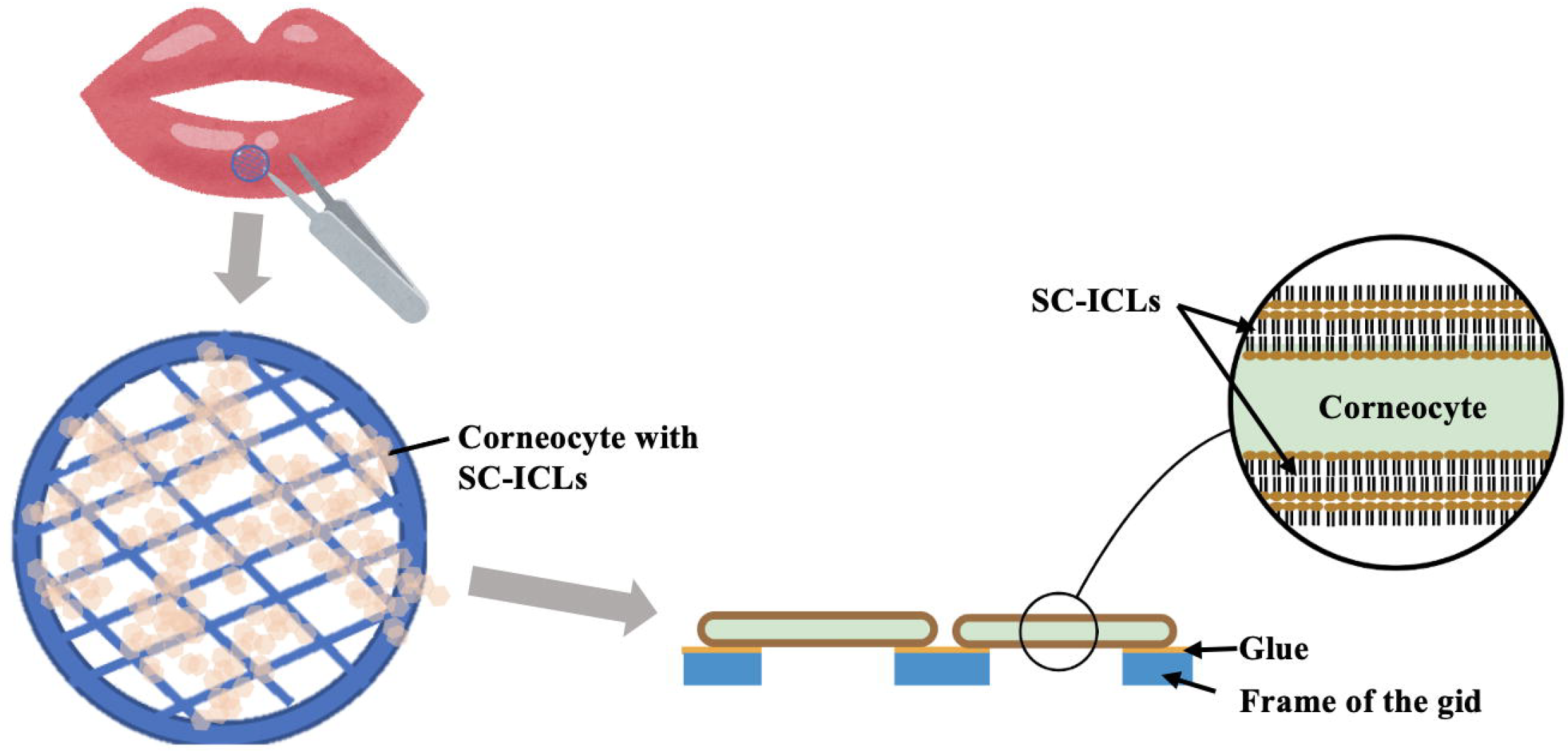
An Illustration of the experimental method for collecting SC-ICLs from lips using the grid stripping method. SC-ISLs can be collected quasi-noninvasively, and the electron beams are irradiated to a corneocyte collected between the frames of the grid [16, 17].

**Figure 2.**
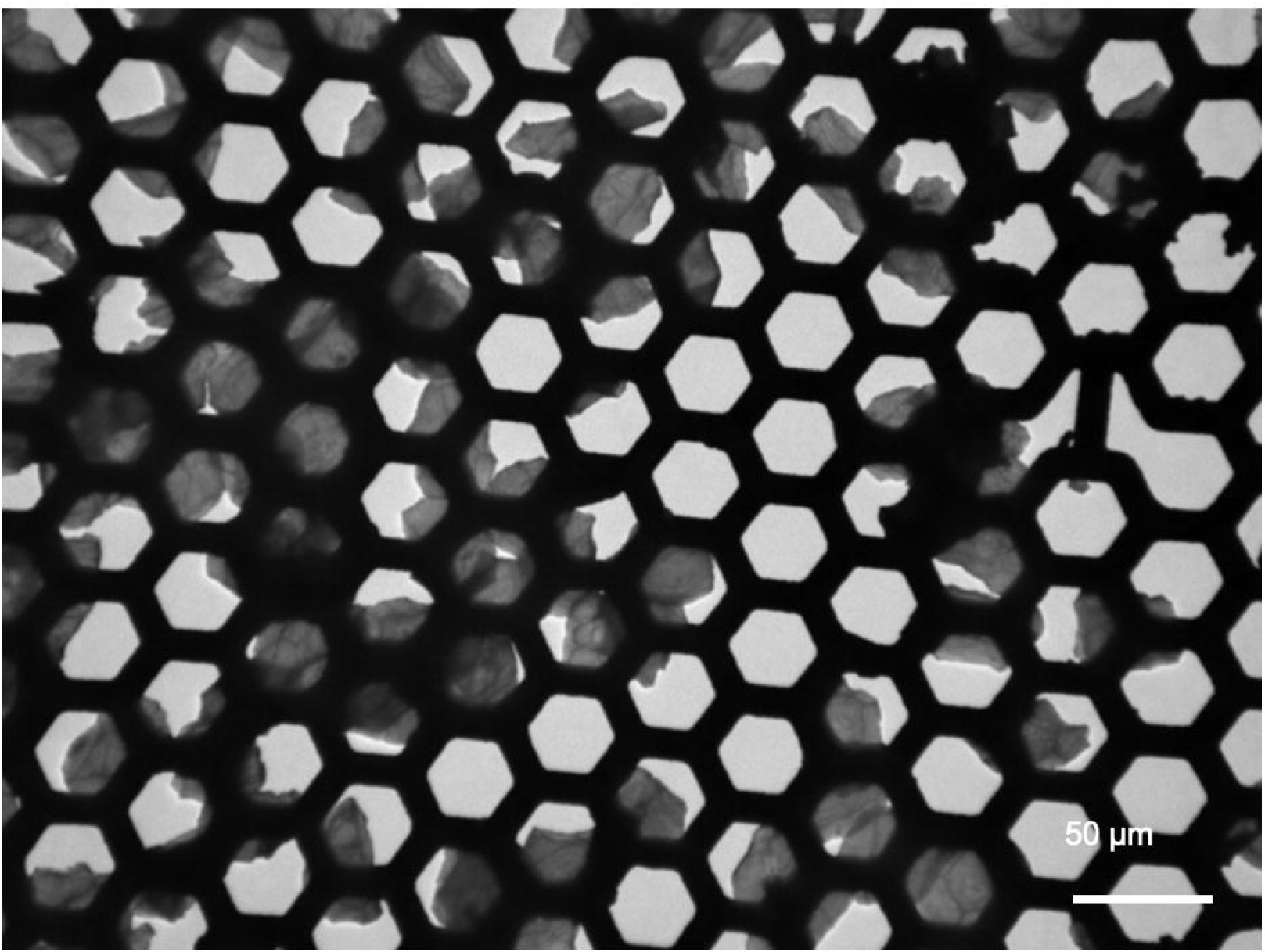
Representative TEM image of lip corneocytes collected by the grid stripping method. The corneocytes (appear gray) were captured on a thin bar hexagonal grid.

**Figure 3.**
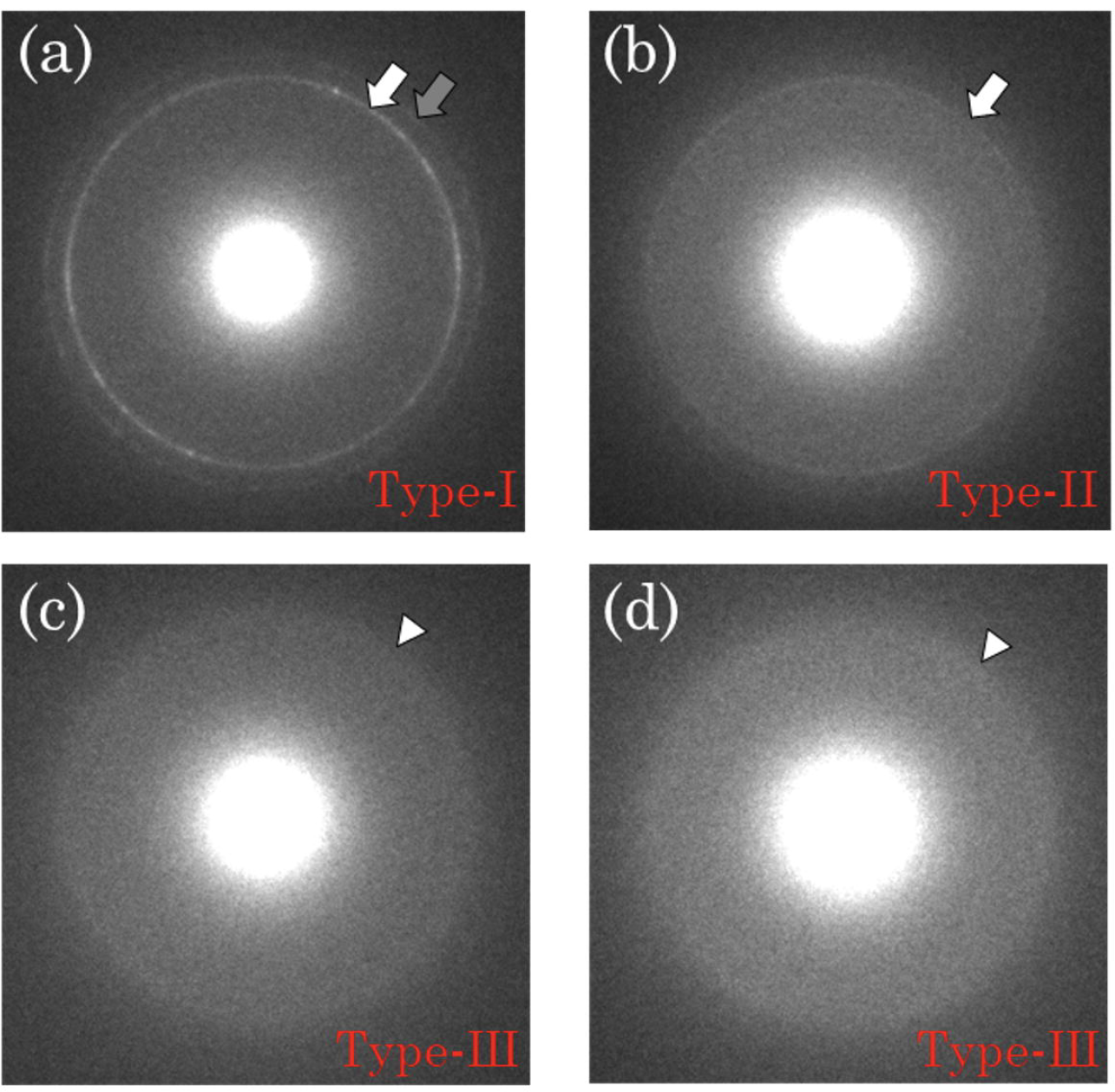
Electron diffraction images (a-d) obtained from lip corneocytes. Type-I: (a) An ED image with two clear Debye-Scherrer rings, one at *s* ≈ 2.4 nm^−1^ (white arrow) and the other at *s* ≈ 2.7 nm^−1^ (grey arrow). Type-II: (b) An ED image with one clear Debye-Scherrer ring at *s* ≈ 2.4 nm^−1^ (white arrow). Type-III: (c and d) ED images with one broad Debye-Scherrer ring at around *s* ≈ 2.2 nm^−1^ (white arrowhead).

**Figure 4.**
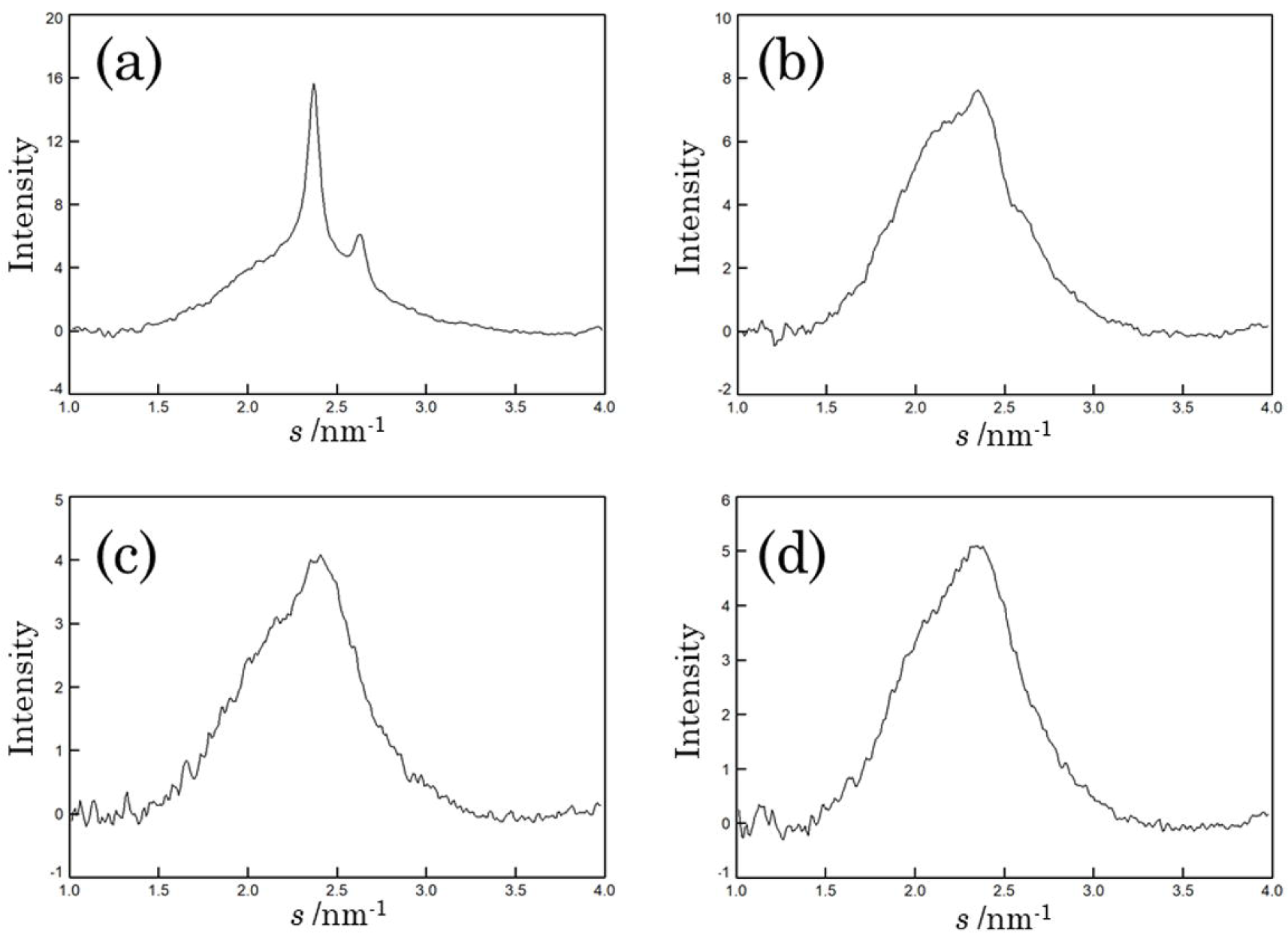
Diffraction intensity profiles (a-d) obtained from lip corneocytes. Type-I: (a) A diffraction intensity profile with two peaks, one at *s* ≈ 2.4 nm^−1^ and the other at *s* ≈ 2.7 nm^−1^. (b) A diffraction intensity profile with a peak at *s* ≈ 2.4 nm^−1^. Type-III: (C and D) ED images with one broad Debye-Scherrer ring at around *s* ≈ 2.2 nm^−1^ (white arrowhead). Type-III: (c and d) Diffraction intensity profiles with a peak at around *s* ≈ 2.2 nm^−1^.

We found no difference in the fundamental characteristics of the diffraction patterns between the above type-I ED images and the previously reported typical ED images of forearm corneocytes [17, 24], which agree well with the corresponding X-ray diffraction images [18, 19]. These results suggest that orthorhombic and hexagonal phases coexist in areas of lip SC-ICLs in a manner similar to those in skin SC-ICLs. The type-II ED image indicating the predominant existence of hexagonal phase is also commonly seen in skin corneocytes, especially in facial sites [17]. These findings demonstrate the existence of highly ordered lipid organization in at least some areas of lip SC-ICLs, similar to skin SC-ICLs.

In contrast, the appearance of type-III ED images is quite rare in the normal forearm skin corneocyte specimens, which usually give sharp rings at 2.4 nm^−1^ and 2.7 nm^−1^. The broad Debye-Scherrer ring at around 2.2 nm^−1^ in the type-III ED image suggests the presence of a fluid phase in the structure of SC-ICLs [14, 24]. However, it is usually hard to evaluate the contribution of the lipid fluid phase to the broad ring at around *s* ≈ 2.2 nm^-1^ because of difficulty in peak deconvolution. As supplemental data, we cooled the specimens to detect the fluid phase and obtained additional experimental evidence suggesting that some cells in lip SC are surrounded by SC-ICLs in the fluid phase (supplemental data 1).

In this study, we employed the LFED method to analyze the lateral packing structure of lip SC-ICLs because it allowed us to collect the sample quasi-noninvasively. Our findings represent the first report on the detailed lateral packing structure of SC-ICLs in lip, suggesting that the LFED method is very useful for evaluating lipid organizations in the SCs, from which non-invasive sample collection is required. It revealed that the lip SC-ICLs had essentially the same structure as those in the skin SC-ICLs, i.e., orthorhombic, hexagonal, and fluid phases, and that the characteristic feature specific to lips is the frequent appearance of the type-III ED images, which suggests extensive existence of the fluid phase.

### Relationship between *A*_*f*_ and lip SC functions

Since it is generally known that the ratio of orthorhombic to hexagonal phase correlates with skin barrier function [15, 24], the lip SC-ICLs structures analyzed in the present study must be also related to the principal lip-SC functions, i.e., barrier and moisturization functions. However, analyzing them using a conventional method faces difficulties due to the extensive distribution of the fluid phase rather than the ordered phases. Estimating the proportion of the fluid phase from diffraction intensity profiles remains challenging, and there is currently no established method for simultaneously assessing both the fluid and ordered phases. Therefore, to clarify the relationship between lip SC-ICL structure and its function, we calculated the frequency of appearance of type-III ED images *A*_*f*_ [%] defined by

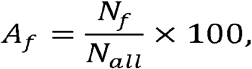

where *N*_*f*_ and *N*_*all*_ are the number of type-III ED images and all the ED images observed for each subject, respectively. The averaged value of *A*_*f*_ for all the subjects was 63% (Table I). The lowest *A*_*f*_ value was 12% (Table I). Preliminary experiments on the arm SC gave the averaged *A*_*f*_ value of 7%. The high value of *A*_*f*_ in lip SC explains well that the barrier function in lip SC is much lower than that in skin SC [9, 10]. Even the lowest *A*_*f*_ value of 12% is higher than the values obtained in normal skin SCs.

**Table I.**
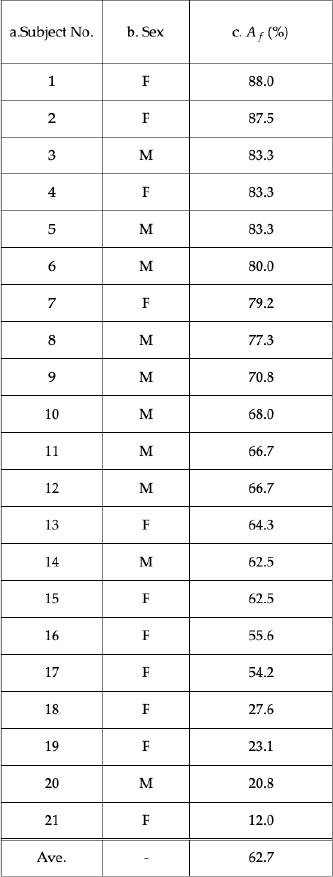
Appearance ratios of ED images with one broad Debye-Scherrer ring at *s* ≈ 2.2nm^−1^ (*A*_*f*_).

| a. Subject No. | b. Sex | c. $A_f$ (%) |
| --- | --- | --- |
| 1 | F | 88.0 |
| 2 | F | 87.5 |
| 3 | M | 83.3 |
| 4 | F | 83.3 |
| 5 | M | 83.3 |
| 6 | M | 80.0 |
| 7 | F | 79.2 |
| 8 | M | 77.3 |
| 9 | M | 70.8 |
| 10 | M | 68.0 |
| 11 | M | 66.7 |
| 12 | M | 66.7 |
| 13 | F | 64.3 |
| 14 | M | 62.5 |
| 15 | F | 62.5 |
| 16 | F | 55.6 |
| 17 | F | 54.2 |
| 18 | F | 27.6 |
| 19 | F | 23.1 |
| 20 | M | 20.8 |
| 21 | F | 12.0 |
| Ave. | - | 62.7 |

We performed a correlation analysis between the structural indicator for lip SC-ICLs, *A*_*f*_ and SC functions, i.e., water content and TEWL. When we used all the examined subjects for the correlation analysis, we found no correlation both between *A*_*f*_ and water content (Fig 5 (a)) and between *A*_*f*_ and TEWL (Fig 5 (b)). Interestingly, we found a fairly clear negative correlation between *A*_*f*_ and water content (Fig 5 (a), correlation coefficient *r* = -0.47, *p* = 0.05) when we picked up the subjects in the group with higher *A*_*f*_ values (> 50%). We consider that subjects with low *A*_*f*_ values should be excluded to clarify the effect of the fluid phase on water content because they seemed to form a separate group where the ordered phases rather than the fluid phase may play a crucial role. It should be noted that the correlation analysis described below represents a unique situation specific to lip SC, where SC-ICL in fluid phase are widely distributed.

**Figure 5.**
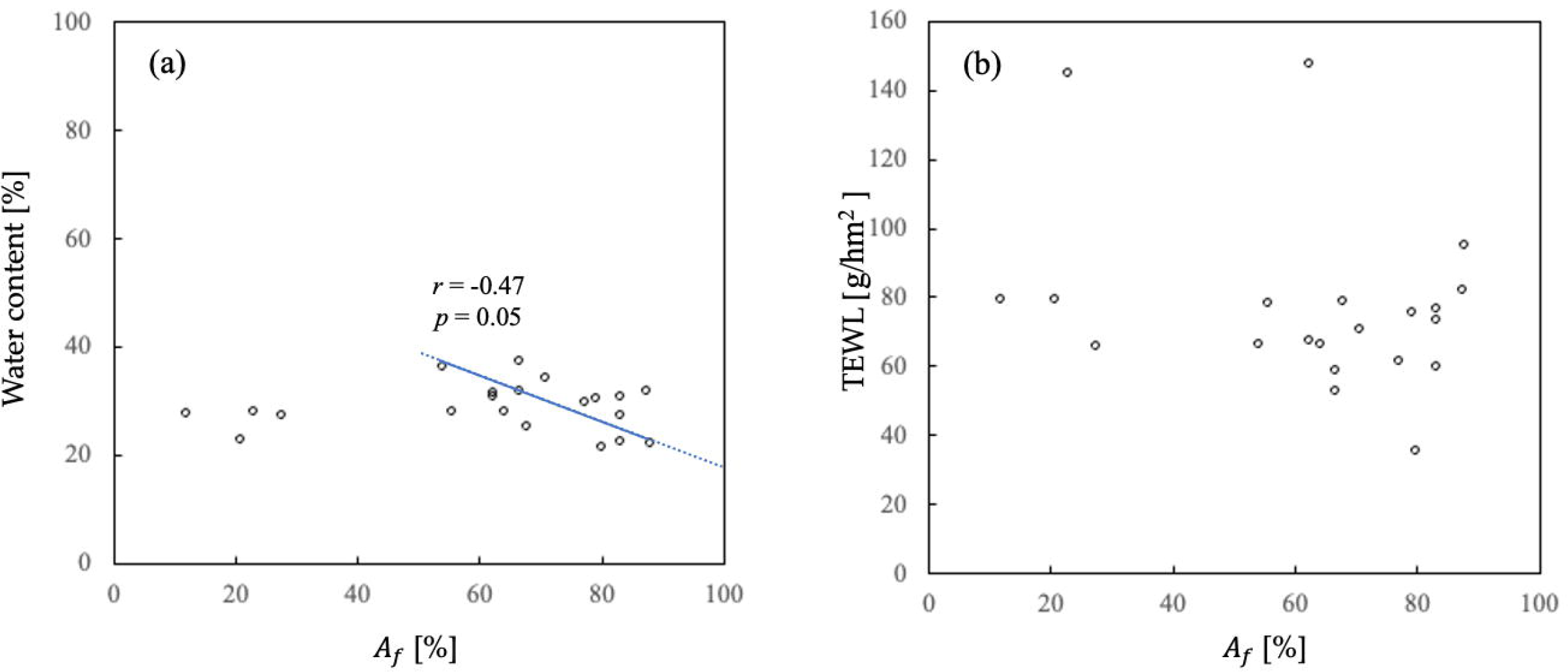
Results of correlation analysis between (a) *A*_*I*_ and water content, and (b) *A*_*f*_ and TEWL. (a): Scatter plot showing no significant correlation between *A*_*f*_ and water content (correlation coefficient *r* = 0.05, *p* = 0.81). However, a negative correlation was observed when picked up the subjects with higher *A*_*f*_ values (*A*_*f*_ > 50%, correlation coefficient *r* = -0.47, *p* = 0.05). (b): No correlation between *A*_*f*_and TEWL (correlation coefficient *r* = -0.29, *p* = 0.20). The standard deviations of *A*_*f*_, water content and TEWL were 23%, 4% and 26 g/hm^2^, respectively.

We evaluated the linear relationship between *A*_*f*_ and water content using standard major axis regression analysis (Fig 5 (a), straight line). The slope was 0.42 and the intercept was 60, i.e., water content [%] *=* -0.42 × *A*_*f*_ + 60. If this linear relationship is valid, its simplest interpretation is to assume the coexistence of two types of cells with a constant volume: one is a corneocyte with a high water content of 60% and the other is an corneocyte with a low water content of (60-*100×0.42=18%*. According to this assumption, the linear dependence of the water content in the whole lip SC on *A*_*f*_ is easily derived. This interpretation, although oversimplified, may be helpful in understanding what determines the water content in lip SC. Moreover, the low water content in lip corneocytes is consistent with the previously reported insufficient formation of natural moisturizing factors (NMF) in lips [25]. If these inferences are correct, the structures/components inside rather than outside the corneocyte are possibly the dominant factor for water content in lip SC.

Obviously, the low water content for the group with *A*_*f*_ values less than 30% cannot be explained in the framework discussed above. In this group contribution of other factor(s) must be taken into account; for example, the relatively low water permeability in the ICLs with ordered phases may restrict the water infiltration into the corneocyte. Further study is required to understand the behavior of water content in lip SC over the whole *A*_*f*_ values. Developing new methods for accurate evaluation of the distribution of ordered and fluid phases, or increasing the number of subjects analyzed, holds the potential to address this problem.

Lastly, we discuss the difference between dependences of water content and TEWL on *A*_*f*_. In contrast to water content, TEWL in lip SC showed no correlation with *A*_*f*_ (Fig 5 (b)). Given that water content is related to the total amount of water in a unit volume of SC, it seems reasonable to assume that the contribution of each cell on water content is additive, as discussed above. In contrast, the contribution of each cell to TEWL may be non-additive because the key factor for TEWL is the continuity of the whole intercellular lipid system, not the properties of each cell. Therefore, it may be possible that the presence of more than a certain percentage of cells with deficient barrier function results in a high saturated value of TEWL, showing no dependence on *A*_*f*_.

## Conclusions

This is an initial report on the application of the LFED method [16-20] for studying the lateral packing structure of human lip SC-ICL. Samples were collected from 21 participants. We found that lip corneocytes showed ED images resembling those of skin SC, suggesting coexistence of intercellular lipids in the hexagonal, orthorhombic and fluid phases. Notably, ED images with no sharp peak (referred to as type-III ED images), which indicate the absence of domains with ordered lipid packing, appeared much more frequently in lip SC than skin SC. Furthermore, the weak intensity of the broad peak at around *s* ≈ 2.2 nm−1 in the type-III ED image suggests a prevalence of corneocytes with thin, fluid phase SC-ICLs in the lip. These results are consistent with the fact that lip has features intermediate between facial skin and oral mucosa [2]. Thus, the LFED method has a great advantage in being able to quasi-noninvasively collect specimens from many subjects and consequently study the lip SC-ICL structures statistically.

Our finding that the fluid phase is predominant in the lip SC-ICLs can qualitatively explain the observed lower water content and higher TEWL than the skin. To further clarify the relationship between lip SC-ICL structure and SC function, we calculated the frequency of appearance of type-III ED images, *A*_*f*_, and analyzed its correlation with water content and TEWL. The analysis revealed a negative correlation between *A*_*f*_and water content in lip SC, particularly when *A*_*f*_ exceeded 50%. If this correlation is correct, it suggests that in the lips with high *A*_*f*_ values (*A*_*f*_ > 50%), the water content is lower in corneocytes surrounded by ICLs in the fluid phase than those surrounded by ICLs in the ordered phases. However, it is important to note that further studies with a larger number of samples are needed to draw a definitive conclusion. Additionally, it is crucial to develop a method that can assess both the fluid and ordered phases simultaneously. Incidentally, we found no correlation between *A*_*f*_ and TEWL possibly because TEWL reached the uppermost value irrespective of *A*_*f*_ owing to a high permeability path composed of the fluid phase.

Finally, the above results suggest that the ratio of ICLs giving type-III ED images is a determinant factor for the water content in lip SC, and this may have implications for chapped lips. The frequent appearance of ICLs with low crystallinity may cause chapping of the lips triggered by dehydration of the lip SC, since water content is negatively correlated with lip roughness [11, 12]. This may give a clue to improving the treatment of chapped lips with conventional lip care products, which contain only occlusive oils or waxes [26]. For example, formulations that can deliver ordered-phase-inducing lipids such as long-chain ceramides [27] into lip SC would be effective for the treatment of chapped lips. Clarifying the structural basis for chapped lips will be the next target studied by our group using the LFED method.

## References

1. Stephen ID, McKeegan AM. Lip colour affects perceived sex typicality and attractiveness of human faces. Perception. 2010;39(8):1104–10.

2. Peramo A, Marcelo CL, Feinberg SE. Tissue engineering of lips and muco-cutaneous junctions: in vitro development of tissue engineered constructs of oral mucosa and skin for lip reconstruction. Tissue Eng Part C Methods. 2012;18(4):273–82.

3. Eroschenko VP. Atlas of Histology with Functional and Clinical Correlations, 12th ed. pp. 285–287, Philadelphia: Lippincott Williams & Wolters Kluwer business; 2013.

4. Xian ZY, Suetake T, Tagami H. Number of cell layers of the stratum corneum in normal skin-relationship to the anatomival location on the body, age, sex and physical parameter. Arch. Dermatol. Res., 1999;291(10):555–9.

5. Hashimoto K. Fine structure of horny cells of the vermilion border of the lip compared with skin. Archs. Oral Biol., 1971;16(4):397–410.

6. Tamura E, Ishikawa J, Naoe A, Yamamoto T. The roughness of lip skin is related to the ceramide profile in the stratum corneum. Int J Cosmet Sci. 2016;38(6):615–21.

7. Moore DJ, Rawlings AV. The chemistry, function and (patho)physiology of stratum corneum barrier ceramides. Int J Cosmet Sci. 2017;39(4):366–72.

8. Ishikawa J, Shimotoyodome Y, Ito S, Miyauchi Y, Fujimura T, Kitahara T, Hase T. Variations in the ceramide profile in different seasons and regions of the body contribute to stratum corneum functions. Arch. Dermatol. Res., 2013;305(2):151–62.

9. Kobayashi H, Tagami H. Functional properties of the surface of the vermilion border of the lips are distinct from those of the facial skin. Br. J. Dermatol., 2004;150(3):563–7.

10. Tagami H. Location-related differences in structure and function of the stratum corneum with special emphasis on those of the facial skin. Int J Cosmet Sci. 2008;30(6):413–34.

11. Arai S, Oshida K, Hikima T, Hasunuma K. Study on lip surface — Characteristics of chapped lip —. J. Jpn. Cosmet. Sci., 1989;13(2):64–68.

12. Kim J, Yeo H, Kim T, Jeong ET, Lim JM, Park SG. Relationship between lip skin biophysical and biochemical characteristics with corneocyte unevenness ratio as a new parameter to assess the severity of lip scaling. Int J Cosmet Sci. 2021;43(3):275–282.

13. Hikima R, Igarashi S, Ikeda N, Matsumoto M, Hanyama A, Egawa Y, et al. Development of lip treatment on the basis of desquamation mechanism. Int J Cosmet Sci. 2004;26(3):165.

14. Hatta I. Stratum Corneum Structure and Function Studied by X-ray Diffraction. Dermato. 2022;2(3):79–108.

15. Bouwstra JA, Ponec M. The skin barrier in healthy and diseased state. Biochim Biophys Acta. 2006;1758(12):2080–95.

16. Nakazawa H, Imai T, Hatta I, Sakai S, Inoue S, Kato S. Low-flux electron diffraction study for the intercellular lipid organization on a human corneocyte. Biochim Biophys Acta. 2013;1828(6):1424–31.

17. Nakazawa H, Imai T, Hatta I, Kato S. Low-flux electron diffraction study on body site dependence of stratum corneum structures in human skin. Biochim Biophys Acta Biomembr. 2022;1864(9):183933.

18. Imai T, Nakazawa H, Kato S. Thermal phase transition behavior of lipid layers on a single human corneocyte cell. Chem Phys Lipids. 2013;174:24–31.

19. Pilgram GS, Pelt AME, Bouwstra JA, Koerten HK. Electron Diffraction Provides New Information on Human Stratum Corneum Lipid Organization Studied in Relation to Depth and Temperature. J. Invest. Dermal., 1999;113(3):403–9.

20. Pilgram GS, Pelt AME, Bouwstra JA, Koerten, Cryo-electron diffraction as a tool to study local variations in the lipid organization of human stratum corneum. J. Microsc., 1998;189(1):71–8.

21. Choe C, Lademann J, Darvin ME. A depth-dependent profile of the lipid conformation and lateral packing order of the stratum corneum in vivo measured using Raman microscopy. Analyst. 2016;141(6):1981–7.

22. Chong Y, Dong R, Liu X, Wang X, Yu N, Long X. Stereophotogrammetry to reveal age-related changes of labial morphology among Chinese women aging from 20 to 60. Skin Res Technol. 2021;27(1):41–8.

23. Yasumori H, Tamura E, Tsukahara K, Inoue Y, Yamamoto T. Age-Related Changes in Lip Morphological Characteristics in Japanese Women. Journal of Society of Cosmetic Chemists of Japan. 2019;53(4):287–96.

24. Pilgram GS, Vissers DC, van der Meulen H, Pavel S, Lavrijsen SP, Bouwstra JA, et al. Aberrant lipid organization in stratum corneum of patients with atopic dermatitis and lamellar ichthyosis. J Invest Dermatol. 2001;117(3):710–7.

25. Bielfeldt S, Laing S, Sadowski T, Gunt H, Wilhelm KP. Characterization and validation of an in vivo confocal Raman spectroscopy led tri-method approach in the evaluation of the lip barrier. Skin Res Technol. 2020;26(3):390–7.

26. Tamura E, Yasumori H, Yamamoto T. The efficacy of a highly occlusive formulation for dry lips. Int J Cosmet Sci. 2020;42(1):46–52.

27. Ohnari H, Naru E, Ogura T, Sakata O, Obata Y. Phase Separation in Lipid Lamellae Result from Ceramide Conformations and Lateral Packing Structure. Chem. Pharm. Bull., 2021;69:72–80.

